# Structural insights into interdomain interactions in *Entamoeba histolytica* APS kinase

**DOI:** 10.64898/2026.04.02.716029

**Authors:** Ryo Hatanaka, Yukiko Ohsumi, Hiroki Matsui, Ayuna Inoguchi, Hina Yuasa, Fumika Mi-ichi, Jun-ichi Kishikawa, Tomoo Shiba

## Abstract

The biosynthetic pathway of 3′-phosphoadenosine-5′-phosphosulfate (PAPS) is a universal and essential metabolic process in many organisms, providing the activated sulfate donor required for the synthesis of diverse sulfated metabolites. However, this pathway has undergone substantial evolutionary diversification among species. In *Entamoeba histolytica*, PAPS biosynthesis occurs within the mitosomes, mitochondrion-related organelles (MROs), representing a distinctive example of lineage-specific evolutionary adaptation. PAPS synthesis proceeds through a conserved two-step, which is sequentially catalyzed by ATP sulfurylase (AS) and adenosine 5′-phosphosulfate (APS) kinase (APSK). In this study, we focused on *E. histolytica* APSK (*Eh*APSK). *Eh*APSK contains an additional AS-like domain (SLD), although its functional role remains unclear.

Here, we determined the crystal structure of full-length *Eh*APSK at 2.60 Å resolution and the structure of the truncated *Eh*APSK lacking APS kinase domain (KD) (*Eh*APSK_ΔKD_) at 2.10 Å resolution. Structural analyses revealed that the SLD engages in dynamic contacts with the KD. Furthermore, deletion of the domain and mutational analyses indicated that the SLD significantly influences the catalytic activity of the KD. Based on these findings, we propose a new regulatory mechanism in which transient interdomain interactions modulate APS kinase activity, representing an unique evolutionary adaptation of *E. histolytica*.

## Introduction

Sulfation of biomolecules is an essential biochemical modification that plays critical roles in numerous physiological processes, including hormone regulation, secretion of signaling molecules, skeletal development, detoxification of xenobiotics, and various forms of cellular proliferation and differentiation, thereby contributing fundamentally to the maintenance of life in living organisms [1,2,3,4]. However, a sulfate is chemically stable and relatively unreactive and therefore must first be converted into an activated form before it can be utilized in sulfation reactions [4,5,6,7]. This sulfate activation pathway is a highly conserved metabolic pathway found across bacteria, protists, fungi, plants, and metazoans.

In sulfate activation pathway, 3’-phosphoadenosine-5’-phosphosulfate (PAPS), the universal sulfate donor, is synthesized through two sequential enzymatic reactions catalyzed by ATP sulfurylase (AS) and adenosine 5’-phosphosulfate kinase (APSK). In the first step, AS catalyzes the synthesis of adenosine 5’-phosphosulfate (APS) and pyrophosphate (PP*i*) from ATP and sulfate ion. In the second step, APS is phosphorylated using ATP by APSK to produce PAPS. After activation pathway, the PAPS serves as a sulfate donor for sulfotransferases involved in the modification of carbohydrates, proteins, lipids, glycolipids, and small organic molecules [7,8,9,10]. In plants and eubacteria, APS and PAPS also function as intermediates in sulfate assimilation pathways responsible for the synthesis of sulfur-containing amino acids such as cysteine and methionine, as well as essential sulfur-containing cofactors including Fe–S clusters and vitamins [4,11,12,13,14,15]. Previous studies have suggested that the APS phosphorylation step catalyzed by APSK constitutes the rate-limiting step in PAPS biosynthesis [16,17,18,19].

A critical biochemical constraint in sulfate activation pathway is that the APS synthesis is intrinsically endergonic and thermodynamically favors the reverse reaction [20,21]. To overcome this barrier, organisms primarily adopt a strategy that biases the reaction by rapidly consuming the intermediates PP*i* and APS through IPP and APSK respectively, thereby maintaining low metabolite concentrations surrounding the pathway. During evolution, these enzymes have diversified into multiple structural arrangements. In higher eukaryotes such as humans and insects, AS and APSK are genetically fused to form bifunctional PAPS synthase (PAPSS). Moreover, trifunctional enzymes containing IPP have been identified in several stramenopile genomes [22]. In some cases, domain fusion not only increases catalytic efficiency but also introduces regulatory functions. For example, in AS from filamentous fungus *Penicillium chrysogenum* (*Pc*), the fused APSK domain lacks APS kinase activity and instead acts as an allosteric regulator of the AS domain [23,24,25].

Among organisms utilizing sulfate activation pathway, the pathogenic facultative anaerobic protozoan *Entamoeba histolytica* show a unique feature. *E. histolytica* is the causative agent of amoebiasis, a parasitic disease that infects approximately 50 million individuals worldwide each year and causes approximately 100,000 deaths annually. Because many infections remain asymptomatic yet transmissible, amoebiasis represents a significant global public health concern [26]. However, the drugs currently available are limited, and no effective vaccine exists. Therefore, the urgent development of new therapeutics against amoebic dysentery is required [27].

*E. histolytica* alternates between an infectious cyst stage and a proliferative trophozoite stage. Both cyst formation and trophozoite proliferation are essential for parasite transmission and pathogenesis. Sulfolipids play key roles in these processes, with cholesteryl sulfate required for cyst formation and fatty alcohol disulfates promoting trophozoite proliferation. The synthesis of sulfolipids relies on PAPS produced by sulfate activation pathway in *E. histolytica* [28,29,30,31].

The sulfate activation pathway in *E. histolytica* exhibits an unusual intracellular localization. In most organisms, this pathway functions in the cytosol, nucleus, or plastids [7,32,33,34,35]. In contrast, *E. histolytica* localizes the pathway into the mitosomes, a kind of mitochondrion-related organelles (MROs) considered to have arisen through reductive mitochondrial evolution under anaerobic conditions [36]. The mitosomes lack mitochondrial genome and has lost most canonical mitochondrial metabolic pathways, including the tricarboxylic acid cycle, electron transport chain, and fatty acid β-oxidation. Notably, *E. histolytica* locates ATP synthesis and PAPS-consuming pathways outside the mitosomes [36,37,38]. Therefore, the exchange of ATP and PAPS between cytosol and mitosomes are required to maintain the life cycle of *E. histolytica*. The advantage of this seemingly inefficient compartmentalization remains unclear.

APSK of *E. histolytica* is the key enzyme of sulfate activation pathway. In addition to this unprecedented localization, APS kinase from *E. histolytica* (*Eh*APSK) possesses unique structural features that distinguish it from homologs in other organisms. *Eh*APSK appears to be a bifunctional enzyme composed of two domains, AS domain (SD) and APSK domain (KD). However, *Eh*APSK contains an N-terminal AS-like domain (SLD) that is structurally homologous to the SD but lacks catalytic activity, fused to the C-terminal KD [28,36,39]. A similar domain composition has been identified in the chemolithoautotrophic bacterium *Thiobacillus denitrificans* (*Td*), where the SLD is predicted to contribute to stabilization of the hexameric assembly [40,41]. However, the functional role of the SLD in *Eh*APSK remains unknown.

Based on this background, we performed structural and functional analyses of *Eh*APSK, a key enzyme in the sulfate activation pathway of *E. histolytica*. We determined the overall structure of *Eh*APSK at 2.60 Å resolution and the KD-truncated SLD structure (*Eh*APSK_ΔKD_) at 2.10 Å resolution by X-ray crystallography. Combining the mutational analyses, the results further revealed that the SLD not only contributes to oligomer stabilization but also functions as a regulatory domain that enhances catalytic activity through transient interactions with the KD. These findings provide new insights into the catalytic and regulatory mechanisms of *E. histolytica* APS kinase.

## Results

### Overall structure of *Eh*APSK

Diffraction data were collected using crystals obtained from screening. Initially, we attempted phase determination via molecular replacement using the structures of AlphaFold2 predictions or *Td*APSK, which shares high sequence homology; however, these attempts did not yield a clear solution. To overcome this, we employed the single-wavelength anomalous dispersion (SAD) method using a gold complex.

The structure was determined at 2.6 Å resolution by SAD phasing. Structural analysis revealed that the asymmetric unit consists of a tetramer (Fig. S1a, Table S1). In contrast, cryo-electron microscopy (cryo-EM) analysis (Fig. S2a,b) produced a density map indicating a dimer, suggesting that the biological assembly in solution is a dimer (Fig. 1a). These observations indicate that, within the crystal lattice, two W-shaped dimers are arranged face-to-face to form the tetrameric assembly. This W-shaped homodimeric architecture has been previously reported in *Aquifex aeolicus* PAPSS (*Aa*PAPSS), as well as within the hexameric assemblies of *Td*APSK and *Pc*AS, suggesting that it is a conserved structural motif within this enzyme family [25,41,42].

**Figure 1.**
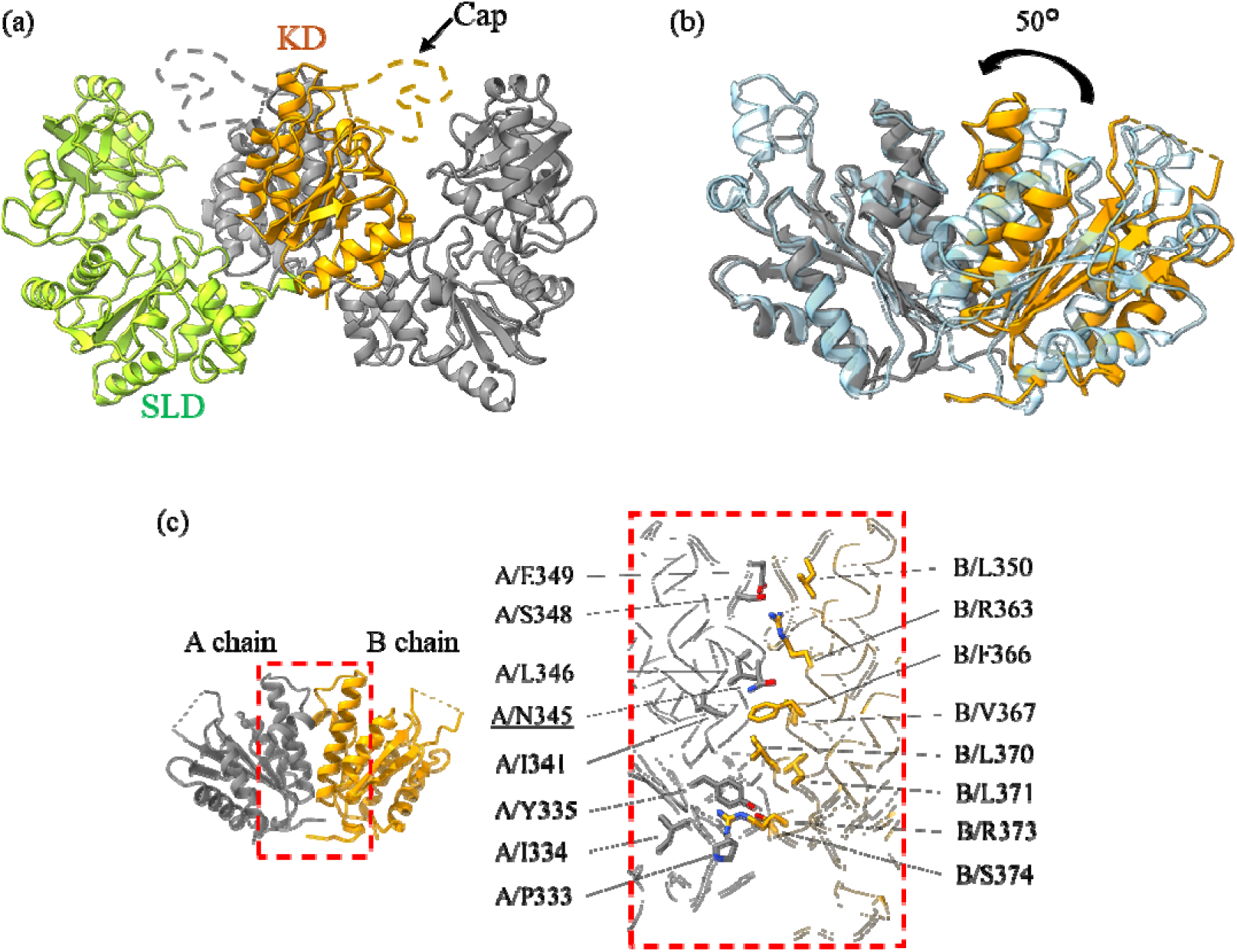
Crystal structure of *Eh*APSK. (a) *Eh*APSK forms a homodimer (colored and gray), and each protomer consists of SLD and KD. In one protomer, SLD and KD are shown in green and orange, respectively, while the cap region of KD could not be modeled and is indicated by orange or gray dashed line. (b) Structural comparison is shownd by superimposing one protomer of *Mt*APSK (cyan, PDB ID: 4BZP) onto the gray protomer of *Eh*APSK. The KD homodimer of *Eh*APSK (orange and gray) is rotated by approximately 50° relative to *Mt*APSK, resulting in a conformation in which the APS-binding sites are positioned closer to each other. (c) Interactions at the KD homodimer interface are shown. Residues involved in intermolecular interactions are shown in stick model, and residues considered important for formation of the “closed” conformation are indicated by underlining. A total of 32 residues participate in interfacial interactions considering both chains A and B.

Each *Eh*APSK protomer is composed of an N-terminal SLD (M1–E293) and a C-terminal KD (L303–K477) (Fig. 1a). structural comparison between the determined EhAPSK structure and the models used for molecular replacement (AlphaFold2 and TdAPSK) reveals a marked difference in the relative orientation and distance between these two domains. This significant domain-level conformational discrepancy likely explains why the initial attempts at phase determination via molecular replacement were unsuccessful. The dimeric structure is primarily formed through interactions at the KD interface, involving residues from both chain A and B, including I334, Y335, N345, E349, R363, F366, L370, and R373 (Fig. 1c), which is generally consistent with many previously reported AS and APSK enzymes possessing the KD or APS kinase-like domain (KLD)[25,41,42]. The KD dimer of *Eh*APSK is tightly packed through the interactions listed above, resulting in the APS-binding sites being positioned close to one another. This compact KD arrangement is also observed in *T. denitrificans* APSK(PDB ID: 3CR8), *Methanothermococcus thermolithotrophicus* APSK (PDB ID: 8A8H), the allosteric domain of *P. chrysogenum* AS (PDB ID: 1I2D), and the low-activity KD of the thermophilic bacterium *A. aeolicus* PAPSS (PDB ID: 2GKS). Hereafter, these are referred to as “closed” KD dimer. In contrast, the APS-binding sites in the KD dimer of *Homo sapiens* PAPSS (PDB ID: 1X6V, 8I1O), *Arabidopsis thaliana* (*At*) APSK (PDB ID: 3UIE), and *Mycobacterium tuberculosis* APSK (PDB ID: 4BZP) exhibit a relatively wider separation (Fig. 1b). These KD dimers are hereafter referred to as “open” KD dimer. Multiple sequence alignment focusing on the KD-corresponding regions of structurally characterized homologs revealed that species adopting the “open” dimer configuration consistently possess glycine at the position corresponding to N345 in *Eh*APSK (Fig. S3). This substitution maybe changes interactions with residues surrounding R363 in *Eh*APSK, thereby contributing to differences in dimer configuration.

### The SLD of *Eh*APSK lost substrate binding capacity

As described above, each protomer consists of two domains: the SLD and the KD. These domains are connected by a short loop (I294–G302). Comparison of the SLD structure with the SDs and the SLD from other organisms showed that the root-mean-square deviation (RMSD) ranged from 0.9 to 1.3 Å (Table S2a), indicating that the overall domain structure, including the Rossmann fold, is well conserved. However, comparison with the SLD of *Aa*PAPSS, which shows the highest sequence identity of the SD (22.6%) among available structures, revealed several distinct structural differences in *Eh*APSK, including shortening of specific α-helices and loops and the presence of an additional N-terminal α-helix (Fig. S4). These features appear to be characteristic of *Eh*APSK.

Previous studies have suggested that *Eh*APSK lacks AS catalytic activity [28,36,39]. The structural analyses in this study suggested two explanations for this loss of activity (Fig. 2). First, the substrate-binding loop that forms part of the catalytic site in catalytically active AS enzymes and contributes to ATP interaction is absent in *Eh*APSK, suggesting intrinsically low affinity for the substrate. Second, the interdomain linker connecting the SLD and the KD is shorter in *Eh*APSK. In catalytically active AS enzymes, this linker extends outward with sufficient length and forms a part of the substrate binding site. In contrast, the shortened linker in *Eh*APSK partially covers the binding pocket, likely restricting substrate access.

**Figure 2.**
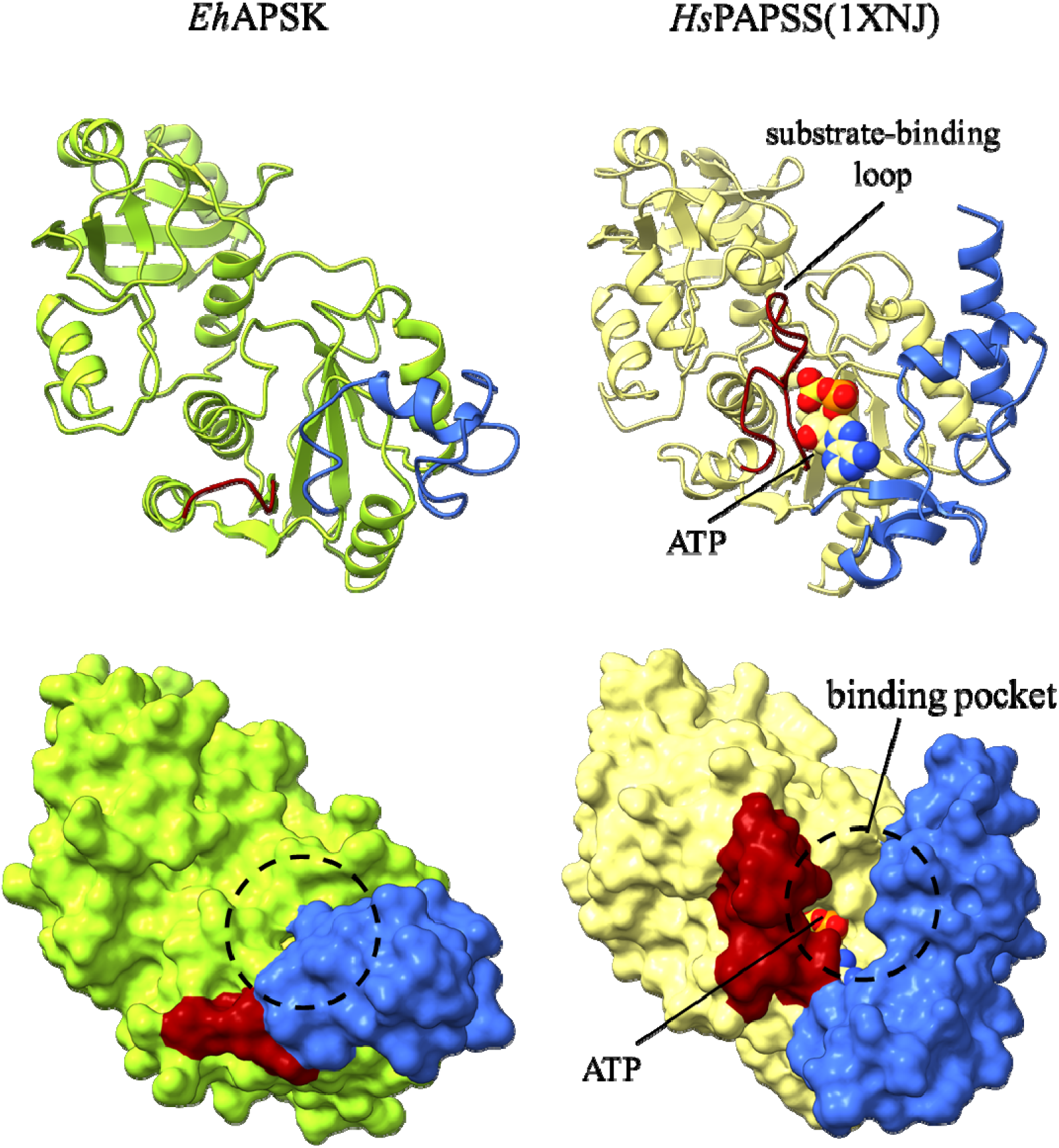
Comparison of SLD in *Eh*APSK and SD. In both panels of the upper row, the loop highlighted in dark red indicates the substrate-binding loop. In *Eh*APSK, the substrate-binding loop is absent compared with SD which is catalytically active structures. In both panels of the lower row, the dashed circles indicate the region corresponding to the binding pocket, whereas the region colored in blue represents the linker connecting to KD. In *Eh*APSK, the entrance to the binding pocket is narrowed by the blue region.

### Catalytic site and the cap region of the KD

Structural comparison of the KD with homologous proteins from other organisms yielded RMSD values of 0.9–1.1 Å (Table S2b), comparable to those observed for the SLD. The KD also shows relatively high sequence identity (37–55%) compared with the SLD (14–23%), supporting a high level of structural conservation. Consistent with this, the fundamental kinase architecture, including the P-loop, is well conserved across species. However, a polar residue implicated in substrate binding in other KDs is replaced by V317 in *Eh*APSK(Fig. S3), which may influence substrate interaction or catalytic activity.

In this study, a spherical electron density was observed at the center of the catalytic site (Fig. 3a). Given that KLSOL was used during crystallization, together with previous reports [43,44,45], this density was modeled as a sulfate ion. The sulfate ion is stabilized via hydrogen bonds with the backbone atoms from G312 to S316 and with the side chain of K315 within the P-loop (Fig. 3b), broadly consistent with interactions reported in previous studies [45]. Comparison with AMP-PNP-bound structures of *At*APSK reported previously [43] suggests that this sulfate-binding site corresponds to the β-phosphate of ATP (Fig. 3c). During the catalysis, this site traps the β-phosphate, thereby positioning the γ-phosphate toward the APS and facilitating the reaction. In the previously reported structure of *Hs*PAPSS, ligand binding suggested to displace the first turn of the α-helix, thereby expanding the P-loop [46]. However, structures lacking the sulfate ion were not obtained in this study. Therefore, it remains unclear whether a similar conformational change occurs in *Eh*APSK, and the potential structural influence of sulfate occupying this position has yet to be elucidated.

**Figure 3.**
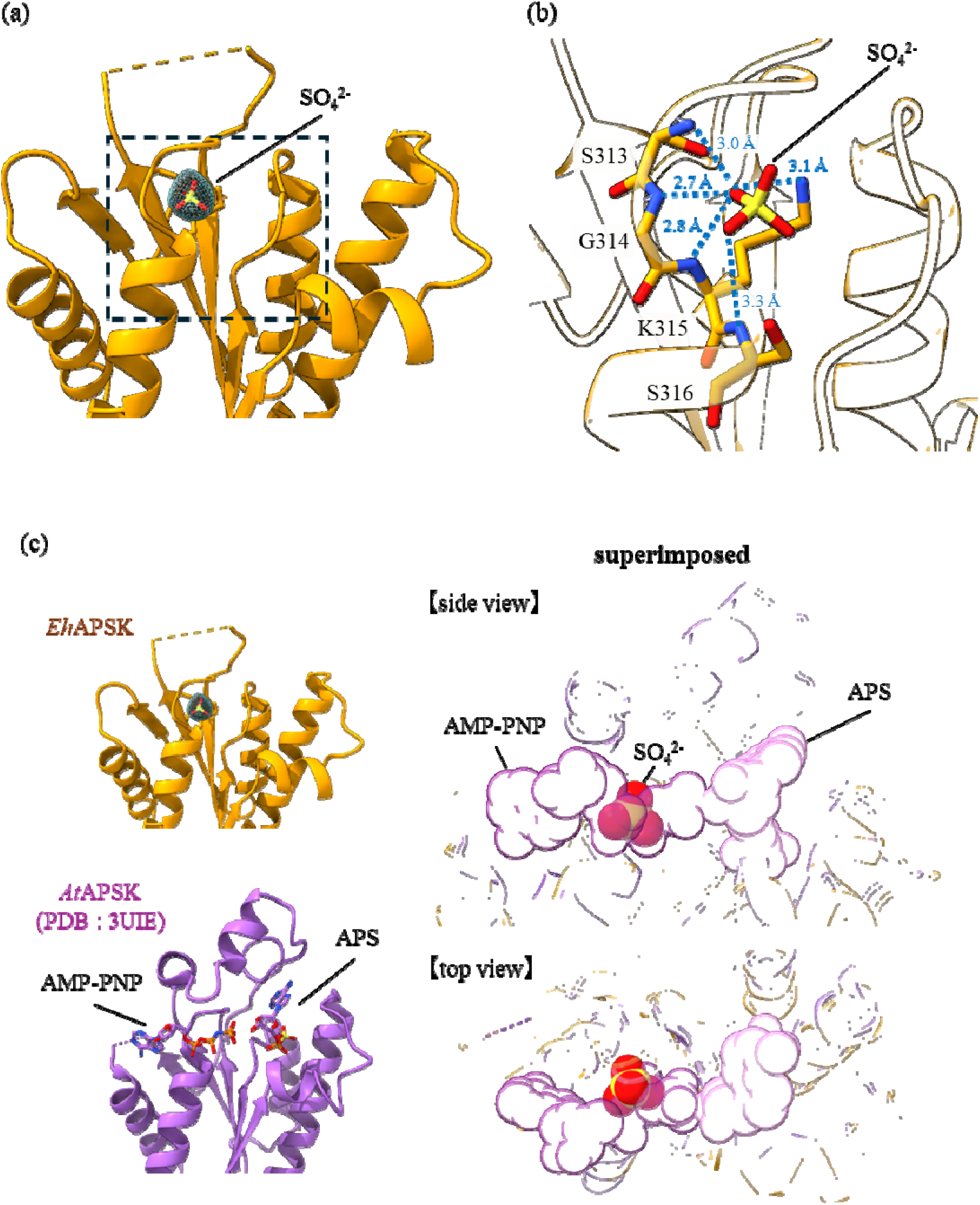
Active site of KD. (a) Spherical electron density observed at the active site of *Eh*APSK. The electron density map is shown at a contour level of 1.5σ. Based on the experimental conditions and previous studies, this density was modeled as a sulfate ion (SOL²L). The fitted sulfate ion is displayed as a stick model. (b) Enlarged view of the region indicated by the black dashed box in (a). Blue dashed lines suggest the possibility of interactions with individual residues in the P-loop. (c) Superposition of *Eh*APSK (orange) with the substrate analog AMP-PNP and APS-bound structure (PDB code:3UIE) (violet). The sulfate ion overlaps with the β-phosphate of AMP-PNP.

The electron density corresponding to the cap region (T410–D440), located over the catalytic site of the KD, was not observed and therefore could not be structured (Fig. 1a). In APSKs from other organisms, this region forms part of the substrate-binding site by acting as a lid that clamps over the substrates. Consistent with previous studies [41,44,47], the cap region contains several residues considered important for substrate binding and catalysis. These residues are conserved in *Eh*APSK, including R418, K421, and F435, and these observations suggest that *Eh*APSK likely employs catalytic mechanisms like those of APSKs from other organisms. Similar observation regarding electron density has been reported in ligand-free structures of *Td*APSK [41] and *Pc*APSK [44], and previous studies have suggested that this region is flexible prior to substrate binding and is stabilized by ligands [44,48]. Accordingly, the lack of observable electron density likely reflects the flexibility of the cap region in *Eh*APSK.

On the other hand, although crystals of *Eh*APSK were obtained in the presence of the substrate APS and the substrate analog AMP-PNP with the aim of stabilizing the flexible cap region, the electron density corresponding to this region remained unobservable in the resulting structure. In the crystal structure determined in this study, the tip of the SLD from a neighboring dimer within the asymmetric unit is positioned close to the APS-binding site of the other dimer (Fig. S1b). This arrangement suggests that the SLD tip may physically occlude the APS-binding site, thereby interfering with substrate binding and preventing stabilization of the cap region.

### Functional role of SLD–KD Interdomain Interactions

While no significant interactions were observed between the SLD and the KD within the same protomer, a robust interaction network centered on E90 and Y133 was identified at the interface between the SLD and the KD across neighboring protomers (Fig. 4a). Especially, the main chain of Y133 forms hydrogen bonds with E324 and R325 of another protomer, while its side chain engages in hydrophobic interactions with V459 of another protomer. In addition, E90 forms a hydrogen bond with Y133 and simultaneously makes hydrophobic contact with V317. These interactions may help constrain the relative positioning of the two domains.

**Figure 4.**
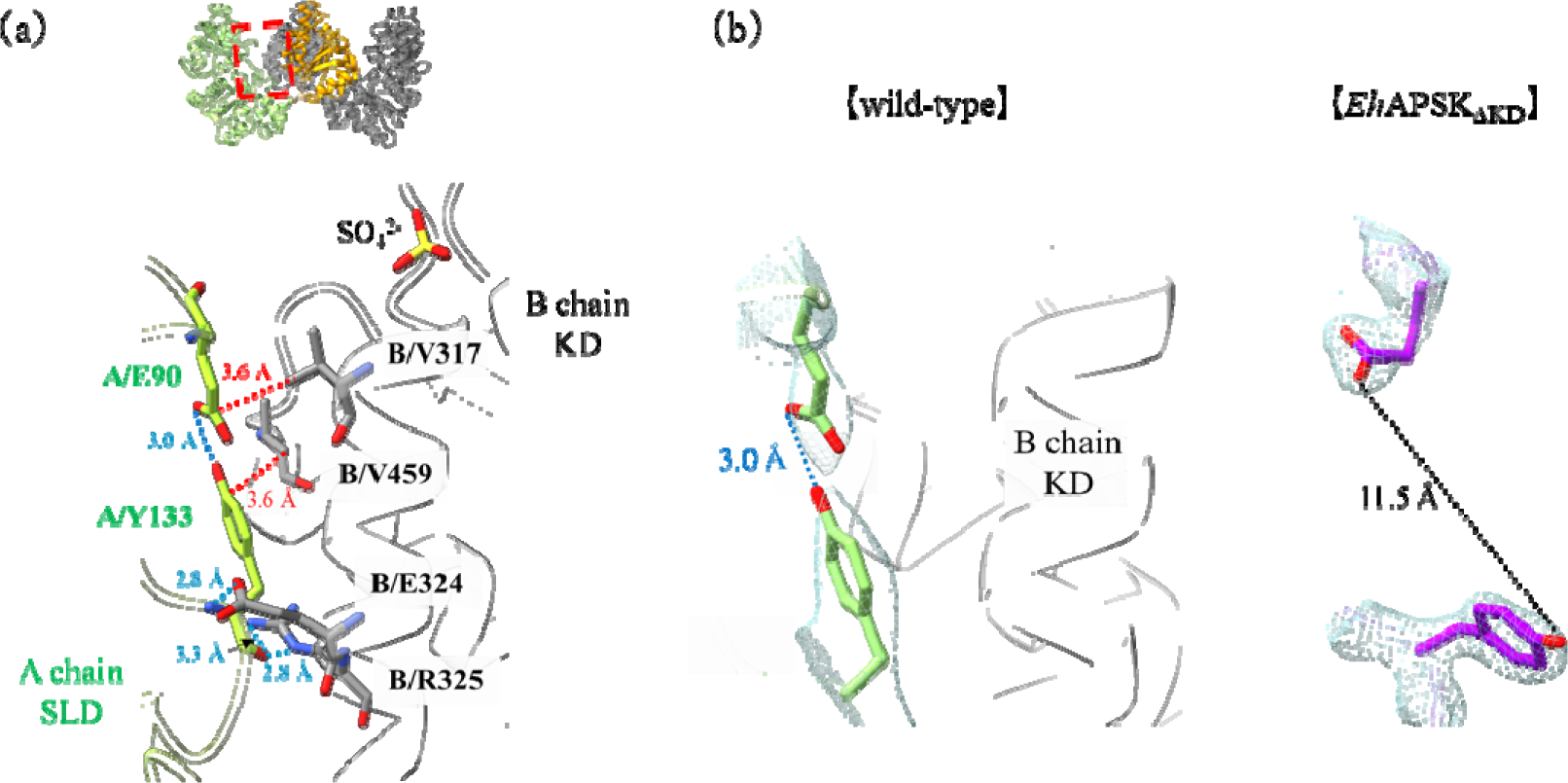
The interaction around E90 and Y133. (a) Interaction network surrounding E90 and Y133. E90 and Y133 in SLD form a complex interaction with the KD. Blue dashed lines indicate hydrogen bonds, and red dashed lines indicate hydrophobic interactions. (b) Comparison of the E90-Y133 pair between wild-type and *Eh*APSK_ΔKD_. The mesh represents electron density maps contoured at 1.0σ for the wild-type and 2.0σ for *Eh*APSK_ΔKD_. In the *Eh*APSK_ΔKD_, black dashed line indicates the altered distance between the two residues. In the absence of the KD, no hydrogen bond was observed between these residues.

To elucidate the functional role of this interaction network between the protomers, the crystal structure of the *Eh*APSK mutant lacking the KD (*Eh*APSK_ΔKD_) was determined at 2.10 Å (Table S1). The RMSD between the SLD of the full-length *Eh*APSK and *Eh*APSK_ΔKD_ was 0.58 Å, indicating that the overall SLD structures are almost identical. However, in *Eh*APSK_ΔKD_, the orientations of E90 and Y133 were changed, and the hydrogen bond between these residues cannot be formed (Fig. 4b). These results suggest that Y133 in the SLD interacts with E324 and R325 in the KD and is coordinated upward, thereby enabling the formation of a hydrogen bond between E90 and Y133. Consequently, interdomain interactions between the SLD and the KD, as described above, are formed. Furthermore, X-ray crystallographic analysis revealed that the *B*-factor, an indicator of molecular flexibility, was markedly reduced in *Eh*APSK_ΔKD_ to approximately 55 Å² compared with the full-length *Eh*APSK (Fig. 5a). This result suggests that the high *B*-factors observed in the full-length *Eh*APSK caused by the flexibility of the interdomain loop and reflect relative domain motions.

**Figure 5.**
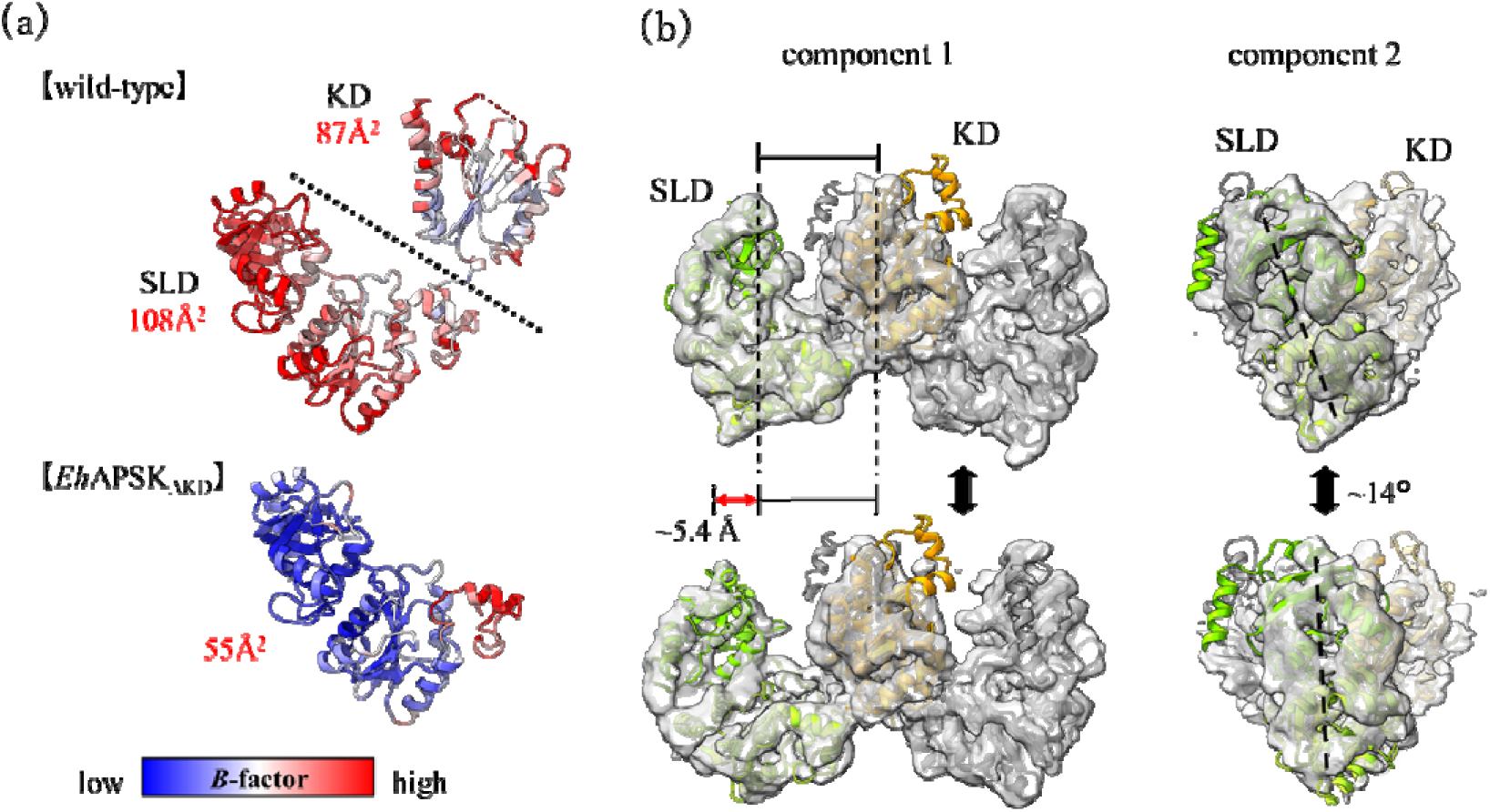
the fructuation of SLD relative to KD. (a) Comparison of *B*-factors between the wild-type *Eh*APSK and *Eh*APSK_ΔKD_. The *B*-factors of *Eh*APSK_ΔKD_ were approximately half of those of the wild-type, indicating reduced structural flexibility. (b) Results of 3D variability analysis from single-particle cryo-EM. Principal component analysis identified two major components describing the flexibility of *Eh*APSK, referred to here as component 1 and component 2. Maps corresponding to the two typical conformations along each component are shown. In addition, the AlphaFold2-predicted structure divided into three segments and these segments were fitted into the maps. The fitted models were used for calculating changes in distance and angle between the KD and SLD. The relative position of these domains varies by approximately ∼5.4 Å and up to ∼14° in rotational displacement.

Supporting evidence for interdomain flexibility was also obtained from cryo-EM analysis. Although single-particle cryo-EM reconstructed a map at a final resolution of 3.44 Å, little or no density was observed for the cap region, consistent with the crystallographic analysis. Additionally, the density corresponding to the SLD was blurry. Subsequent Local refinement and 3D classification in cryoSPARC [49] under various parameters resulted in no significant improvement in the map quality. In contrast, 3D variability analysis in cryoSPARC revealed that the SLD undergoes relative motion with respect to the KD (Fig. 5b). Principal component analysis of the 3D variability identified two major components: one corresponding to the SLD approaching the KD, and the other corresponding to the SLD moving laterally. The relative positions of the SLD and the KD varied by up to approximately 5.4 Å in distance and 14° in angle. The angular and distance changes associated with these motions were estimated by fitting structures predicted by AlphaFold2 [50], generated by dividing *Eh*APSK into three segments: M1-Y133, D134-P297, T298-K477.

Next, we prepared wild-type *Eh*APSK, the *Eh*APSK mutant lacking the SLD (*Eh*APSK_ΔSLD_), and alanine substitution mutants at E90 and Y133, which were predicted to be critical for interdomain interactions. Enzymatic activity assays and kinetic parameter analyses were subsequently performed for each construct.

*Eh*APSK_ΔSLD_ was difficult to express in *Escherichia coli*. To improve expression efficiency, a SUMO tag was introduced at the N-terminal of *Eh*APSK_ΔSLD_, enabling successful expression. However, during purification, the SUMO-cleaved *Eh*APSK_ΔSLD_ exhibited a high tendency to aggregate (Fig. S5a). Although it formed dimers at low concentrations, the protein rapidly became cloudy and precipitated during concentration steps. In contrast, the single-residue mutants (E90A and Y133A) exhibited purification profiles similar to the wild-type *Eh*APSK.

Enzymatic activities of the purified proteins were evaluated by fitting the time-dependent change in A_340_ to the Michaelis–Menten equation (Table 1). Compared with the *k*_cat_ of wild-type (1270 ± 355 s^-1^ for ATP and 1688 ± 244 sL¹ for APS), both *Eh*APSK_ΔSLD_ (9.50 ± 1.03 sL¹ for ATP and 13.8 ± 0.87 sL¹ for APS) and the single-residue mutants (E90A : 21.1 ± 1.12 sL¹ for ATP and 33.8 ± 1.65 sL¹ for APS, Y133A : 59.5 ± 1.78 sL¹ for ATP and 37.1 ± 3.65 sL¹ for APS) exhibited markedly reduced. These values correspond to an approximately two orders-of-magnitude decrease for *Eh*APSK_ΔSLD_ and 21- to 60-fold decreases for the single-residue mutants compared to the wild-type. In contrast, no significant changes were observed in *k*_cat_/*K*_M_ for either substrate. These results indicate that interactions between the SLD and the KD positively regulate catalytic activity.

**Table 1.**
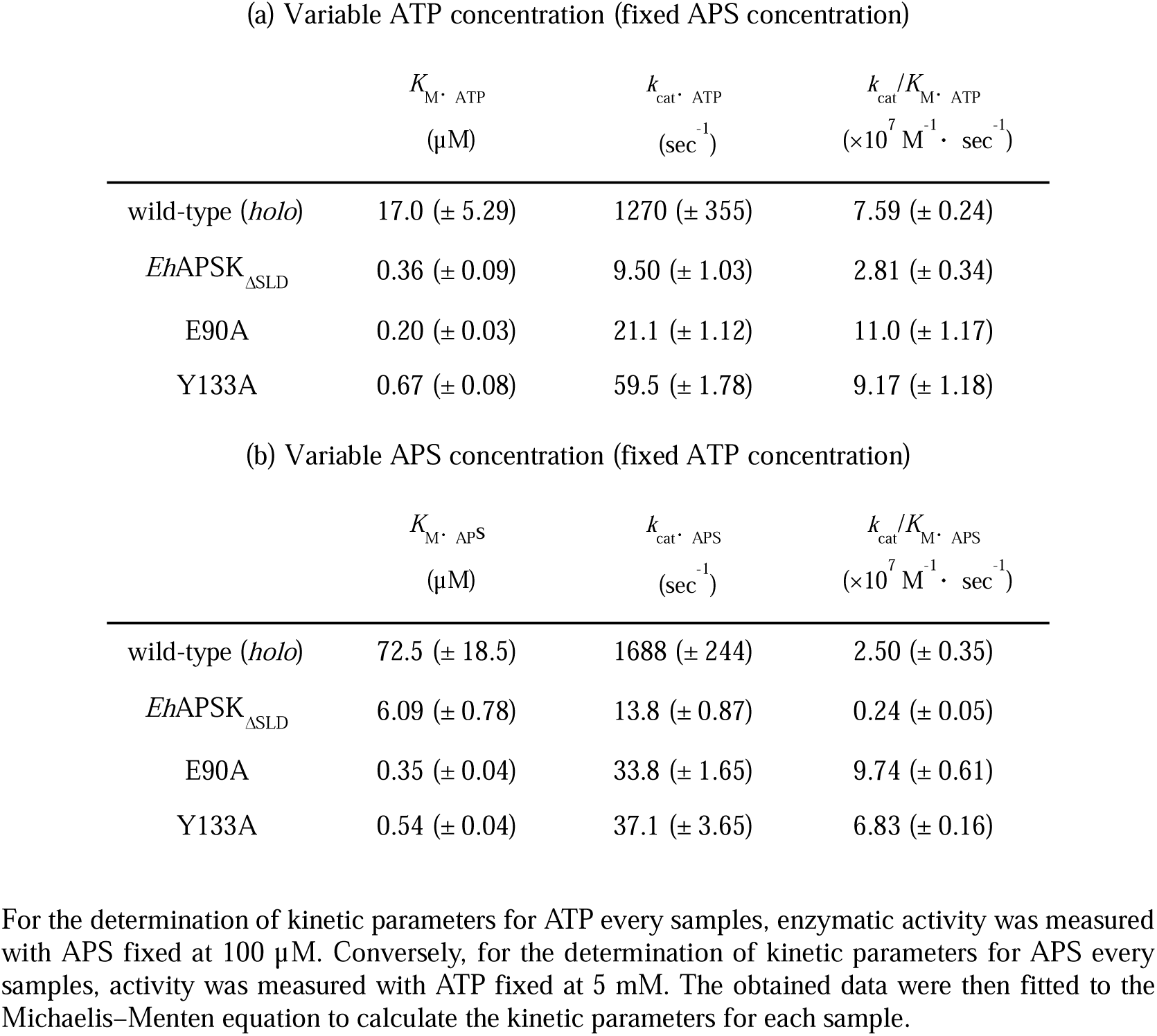
Kinetic parameters of the wild-type enzyme and its mutants.

In *Aa*PAPSS, which possesses structurally similar architectures, Y142 (Fig. S6), corresponding to Y133 in *Eh*APSK, has been suggested to contribute to interdomain interactions [42]; however, no interaction with K80, corresponding to E90 in *Eh*APSK, has been reported. In *Td*APSK, although the residue corresponding to E90 is conserved as E106, the residue corresponding to Y133 is replaced by P142, and no equivalent interaction has been observed [41]. Furthermore, residue pairs analogous to E90 and Y133 are present in *Sc*AS and *Pc*AS; however, these enzymes contain the KLD lacking APSK catalytic activity, suggesting that their catalytic mechanisms differ from that of *Eh*APSK. Taken together, the interdomain interaction identified in this study is a unique structural and functional feature of *Eh*APSK.

## Discussion

In this study, we report the structural and activity analysis of EhAPSK and its mutants, a key enzyme in the sulfate activation pathway of *E. histolytica*. The results provide insight into how EhAPSK mediates the synthesis of PAPS. The uncovered characteristics of *Eh*APSK revealed in this study should reflect the part of the sulfate activation pathway adapted to this unique organellar environment, like mitosome in *E. histolitica*.

In addition, the spherical electron density at the catalytic site of the KD suggests that the ATP phosphate moiety is anchored by the P-loop during catalysis, implying potential competitive inhibition by P*i*. In the sulfate activation pathway, APS and PP*i* are produced by AS, with PP*i* subsequently hydrolyzed to P*i* by IPP, resulting in the accumulation of two P*i* per PAPS in the mitosomes. Accumulated P*i* may inhibit APSK, reducing sulfate activation efficiency. Therefore, maintenance of P*i* homeostasis within the mitosomes is likely required. In this context, the mitosomal phosphate carrier *Eh*PiC [51] is likely essential for sustaining efficient sulfate activation.

The elevated *B*-factor observed in the crystal structure of full-length *Eh*APSK, together with the results of 3D variability analysis in single-particle cryo-EM, indicates that the relative position between the SLD and the KD exhibits substantial conformational flexibility in solution (Fig. 5). In contrast, *Td*APSK, which also contains an SLD, forms a hexamer in solution, where the SLD tightly interacts with the KD of neighboring protomers and the SLD has been proposed to stabilize the hexameric assembly. Therefore, the flexibility between the SLD and the KD in the dimeric *Eh*APSK suggests that its functional role is distinct from that in *Td*APSK. The mutational analyses of *Eh*APSK demonstrate that the SLD in *Eh*APSK enhances kinase activity through the interactions with the KD of the opposite protomer (Table 1).

Integration of these findings suggests the presence of a transient interdomain interaction between the SLD and the KD that positively regulates enzymatic activity. A mechanistic model can be proposed to explain how this transient interaction occurs. In the *apo* state, the SLD fluctuates relative to the KD. When the SLD approaches the KD, the orientation of Y133 is constrained toward E90 through interactions with E324 and R325 in the KD. Subsequently, the hydrogen bond is formed between Y133 and E90, which orients the aliphatic portion of the E90 side chain toward V317, allowing favorable nonpolar contacts that stabilize the SLD–KD interface. AlphaFold2 [50] predicts potential interactions between the tip of the SLD and the cap region of the KD (Fig. S7). This stabilization likely facilitates interactions between residues near the catalytic site, thereby enhancing catalytic activity. Consistent with these observations, such a regulatory mechanism has not been reported in other organisms, suggesting that it represents a novel mechanism of enzymatic regulation first identified in *E. histolytica*. However, the precise interactions by which the SLD influences the catalytic site remain unclear. Therefore, further structural studies of substrate-bound complexes and molecular dynamics simulations will be essential to elucidate the catalytic mechanism in detail.

## Materials & Methods

### Construction of Mutants

The plasmid encoding full-length *Eh*APSK based on the pCold-I™ expression system was used [39]. The pCold-I vector introduces an N-terminal His_6_ tag to the target protein and contains an ampicillin resistance gene as a selection marker.

Using this plasmid as a template, four APSK mutants were constructed in this study: two single-domain truncation mutants corresponding to the SLD and the KD domains, and two point mutants (E90A and Y133A). All mutants were generated using seamless ligation based on homologous recombination according to a previously reported protocol [52]. Primers were designed so that PCR fragments contained homologous sequences at both ends corresponding to adjacent fragments, enabling plasmid assembly through homologous recombination.

PCR amplification was performed using KOD One® PCR Master Mix (Toyobo). The amplified fragments were treated with DpnI (New England Biolabs) to remove template plasmids and subsequently purified using the Gel/PCR DNA Isolation System (VIOGENE). Plasmid construction was then performed using the XE cocktail prepared according to the reported protocol [53].

For *Eh*APSK_ΔSLD_, the pET-SUMO vector was used to improve folding efficiency, allowing expression of a protein containing an N-terminal His_6_-SUMO tag. The pET-SUMO vector contains a kanamycin resistance gene as a selection marker. The plasmid encoding *Eh*APSK_ΔSLD_ was constructed by inserting the corresponding PCR fragment into the pET-SUMO vector using the same seamless cloning procedure [52]. Primer sequences used for mutant construction are listed in supplements (Table S3).

### Protein Expression and Purification

full-length *Eh*APSK, single-domain mutants, and point mutants were expressed in *E. coli* BL21 Star™ (DE3) or NiCo21 (DE3) competent cells, following a previously described protocol [39]. Briefly, 6 mL overnight cultures grown in LB medium containing 100 μg/mL ampicillin or kanamycin were inoculated into 1 L LB medium in 3 L baffled flasks. Cells were cultured at 37□ with shaking until the OD□□□ reached approximately 0.4, followed by cooling in an ice-water bath for 30 min. Protein expression was induced by adding 200 µM isopropyl-β-D-thiogalactopyranoside, and cultures were incubated at 15□ for 17–20 h.

Cells were harvested by centrifugation using a Beckman Avanti J-20 centrifuge equipped with a JLA 8.1000 rotor (8,000 × *g*, 15 min, 4□). Approximately 10 g of cell pellet was resuspended in lysis buffer containing 50 mM Tris-HCl (pH 8.0), 300 mM NaCl, 20 mM imidazole, 10% (w/v) glycerol, 1% Triton X-100, 0.1 mg/mL lysozyme, and 0.25 mM PMSF. After sonication and centrifugation (15,000 × *g*, 30 min, 4□), the supernatant was incubated with Ni-NTA agarose (QIAGEN) pre-equilibrated with equilibration buffer (50 mM Tris-HCl (pH 8.0), 300 mM NaCl, 20 mM imidazole) for 1 h at 4□. The resin was packed into an open column and washed sequentially with 20 column volumes (CV) of equilibration buffer and 10 CV of wash buffer (equilibration buffer + 50 mM imidazole). The target protein was eluted with 10 CV of elution buffer (equilibration buffer + 250 mM imidazole). Eluted fractions were dialyzed overnight at 4□ to remove imidazole and concentrated to approximately 1 mL (10–20 mg/mL) using a 50 kDa cutoff Amicon Ultra-15 concentrator (Millipore).

The concentrated sample was subjected to size-exclusion chromatography (SEC) using an ÄKTA pure system (Cytiva) with an ENrich SEC 650 26/60 HiLoad column equilibrated with SEC buffer (50 mM Tris-HCl (pH 8.0), 300 mM NaCl). Fractions corresponding to the major protein peak were pooled, and high-purity fractions were selected based on SDS-PAGE analysis. Samples were further concentrated to approximately 10 mg/mL, and protein concentration was determined using the molar extinction coefficient (ε□□□ = 25,580 M□¹·cm□¹) calculated by ProtParam (https://web.expasy.org/protparam/). Final samples were mixed with glycerol to a final concentration of 50% and stored at −30□.

For *Eh*APSK_ΔSLD_, the SUMO-tagged KD eluted from the first IMAC step was incubated with SUMO protease at a molar ratio of 400:1 at 4□ for 17–20 h with gentle mixing. The reaction mixture was subjected to a second IMAC step, and the tag-free KD protein was collected in the flow-through and wash fractions which did not contain imidazole. These fractions were concentrated to approximately 1 mg/mL using a 10 kDa cutoff Amicon concentrator and further purified by SEC using an ENrich SEC 70 26/60 HiLoad column. Fractions containing the dimer were pooled and immediately used for activity assays without further concentration. SDS-PAGE results of final purified samples are shown in supplements (Fig. S5b).

### Crystallization

Purified proteins were buffer-exchanged into the crystallization buffer containing 10 mM Tris-HCl (pH 8.0), 100 mM NaCl, and 1 mM TCEP using a 50 kDa cutoff Amicon concentrator.

For full-length *Eh*APSK, the protein was concentrated to approximately 25 mg/mL and incubated on ice for 1 h with 5 mM APS, 5 mM AMP-PNP, and 5 mM MgCl□. Crystallization was performed using the hanging-drop vapor diffusion method at 20□ by mixing equal volumes of protein solution and reservoir solution containing 0.21–0.27 M K□SO□ and 16–18% polyethylene glycol (PEG) 3400. Crystals appeared after approximately one week.

For *Eh*APSK_ΔKD_, the protein was exchanged into the crystallization buffer using a 10 kDa cutoff Amicon concentrator and concentrated to approximately 10 mg/mL. Crystallization was performed using the sitting-drop vapor diffusion method at 20□ by mixing equal volumes of protein solution and PEG/Ion □ Screen™ condition No. 30 (0.2 M ammonium tartrate dibasic (pH 7.0), 20% PEG 3350), yielding crystals after approximately two weeks.

### X-ray Diffraction and Structural Modeling

For phase determination using single-wavelength anomalous dispersion (SAD), crystals of full-length *Eh*APSK were transferred into a cryoprotectant solution containing 1 mM K[Au(CN)□], 30% glucose, 5 mM APS, 5 mM AMP-PNP, 5 mM MgCl□, and reservoir solution with 2% higher PEG 3400 concentration. After soaking for 1–24 h, crystals were flash-cooled in liquid nitrogen.

Native full-length crystals and *Eh*APSK_ΔKD_ crystals were transferred into their respective cryoprotectant solutions and flash-cooled in liquid nitrogen. full-length crystals were transferred into the solution described above without heavy atoms, while *Eh*APSK_ΔKD_ crystals were transferred into PEG/Ion □ Screen™ condition No. 30 supplemented with 25% glycerol.

Diffraction data were collected at beamline BL44XU of SPring-8 (Harima, Japan) and processed using XDS [53]. Initial phases for full-length *Eh*APSK were determined by the SAD method using CRANK2 [54] in CCP4i [55], followed by partial model building with Buccaneer [56,57]. Complete structural modeling was performed using molecular replacement with native datasets in MOLREP [58] or Phaser-MR [59].

The structure of *Eh*APSK_ΔKD_ was determined by molecular replacement using MOLREP, with a search model generated by the removing KD from the full-length structure. Manual model building and crystallographic refinement were iteratively performed using Refmac5 [60] and Coot [61]. The final models have been deposited in Protein Data Bank (PDB ID of full-length *Eh*APSK: 23VK, PDB ID of *Eh*APSK_ΔKD_: 23VL). Structural figures were prepared using PyMOL [62] and UCSF ChimeraX [63].

### Cryo-EM Data Collection and Image Processing

Grid preparation and cryo-EM data collection were performed at the Institute for Protein Research, Osaka University. For grid preparation of full-length *Eh*APSK, SEC peak fractions were diluted to approximately 0.24 mg/mL in SEC buffer and mixed with ligands to final concentrations of 12.5 µM APS, 1.25 mM ATP, and 12.5 mM MgCl□ (*Eh*APSK:ligunds molar ratio = 1:3). A 2.6 μL aliquot of the sample was applied onto glow-discharged UltrAuFoil® R1.2/1.3 (300 mesh) grids and plunge-frozen in liquid ethane using a Vitrobot IV with 3 s blotting time.

Cryo-EM movies were acquired using a Titan Krios transmission electron microscope (Thermo Fisher Scientific) equipped with a K3 BioQuantum camera (Gatan) and automated using SerialEM [64]. Data were collected at 300 kV accelerating voltage, nominal magnification of 105,000, pixel size of 0.675 Å, total dose of 65 e□/Å², defocus range of −0.8 to −1.8 µm, and spherical aberration of 0.066 mm. A total of 11,470 movies were collected.

Image processing workflows are summarized in supplements (Fig. S2a, Table S4). All processing steps were performed using cryoSPARC v4.4.1 [49]. Particle picking was performed using the machine-learning-based Topaz algorithm□□□□. After motion correction and CTF estimation, 1,473 low-quality micrographs were discarded. Automatic picking by Topaz [65] identified 1,923,378 particles, which were extracted and subjected to iterative 2D classification, ab initio reconstruction, and heterogeneous refinement. The final dataset containing 450,150 particles was refined using non-uniform refinement with C1 symmetry and per-particle defocus optimization, yielding a final reconstruction at 3.44 Å resolution. The SLD region exhibited weak and asymmetric density. Local refinement, 3D classification, and 3D variability analysis were performed to further characterize this domain. Map resolution was determined using the gold-standard Fourier shell correlation criterion (FSC = 0.142). Atomic modeling was not performed for the cryo-EM map.

### Activity Assay and Kinetic Analysis

APSK activity of wild-type and mutant proteins was measured following a previously reported protocol [39] using UV-STAR MICROPLATE® (96-well, half-area, clear) plates (Greiner) and a Varioskan ALF plate reader (Thermo Fisher Scientific). Each well contained 180 μL reaction mixture composed of 90 μL two-fold concentrated coupling solution, 6 μL enzyme solution, substrate at varying concentrations, and ultrapure water to adjust the volume.

Reactions were initiated by adding APS or MgATP and monitored by measuring absorbance decrease at 340 nm at 30□ for approximately 1 h. The coupling solution (1×) contained 100 mM Tris-HCl (pH 8.0), 1 mM MgCl□, 1 mM KCl, 1 mM NADH, 1 mM phosphoenolpyruvate, 1 U nucleoside diphosphate kinase, 3.5 U pyruvate kinase, and 5.0 U lactate dehydrogenase. Depending on the assay design, the second substrate was supplied at saturating concentration (5 mM MgATP or 100 µM APS). Blank reactions containing SEC buffer instead of enzyme solution were included, and baseline values were subtracted from all measurements. Initial reaction velocities were determined by linear regression of the initial linear portion of NADH absorbance decrease at 340 nm, recorded at 15–30 s intervals, and converted to reaction rates using the molar extinction coefficient of NADH. The initial velocity data were fitted to the Michaelis–Menten equation:

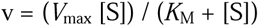

Kinetic parameters including *K*_M_ and *V*_max_ were obtained from the fitted curves. The catalytic turnover number (*k*_cat_) was calculated from *V*_max_ using enzyme concentrations determined spectrophotometrically, and catalytic efficiency was expressed as *k*_cat_/*K*_M_. All measurements were performed at least in triplicate.

## CRediT authorship contribution statement

**Ryo Hatanaka:** Conceptualization, Investigation, Methodology, Visualization, Writing – original draft**. Yukiko Ohsumi:** Investigation**. Hiroki Matsui:** Investigation**. Ayuna Inoguchi:** Investigation**. Hina Yuasa:** Investigation**. Fumika Mi-ichi:** Funding acquisition, Resources, Writing – review and editing**. Jun-ichi Kishikawa:** Conceptualization, Funding acquisition, Methodology, Resources, Supervision, Writing – review and editing**, Tomoo Shiba:** Conceptualization, Funding acquisition, Methodology, Resources, Supervision, Writing – review and editing.

## Acknowledgement

We thank all the staff members of beamline BL44XU (SPring-8) for assistance with X-ray diffraction data collection. The synchrotron radiation experiments were performed at BL44XU of SPring-8 (proposal nos. 2019A6918, 2019B6918, 2020A6518, 2020B6518, 2021A6616, 2021B6616, 2022A6714, 2022B6714, 2023A6814, 2023B6814). The synchrotron beamline BL44XU at SPring-8 was used under the Cooperative Research Program of the Institute for Protein Research, Osaka University. We are thankful to Prof. Takayuki Kato, Mika Hirose and Misaki Arie (Institute of Protein Research) for cryo-EM data collection. Our research was supported by Grant -in -Aid for Scientific Research (JSPS KAKENHI) Grant Numbers 24K10217 to T.S., 25K01949 to J.K. and 25K02488 to F.M. Our research was also supported by the Platform Project for Supporting Drug Discovery and Life Science Research (Basis for Supporting Innovative Drug Discovery and Life Science Research (BINDS)) from AMED under Grant Number JP25ama121001.

## Declaration of generative AI and AI-assisted technologies in the manuscript preparation process

During the preparation of this manuscript, the author used Google Gemini in order to improve the quality of the language. After using this service, the author subsequently reviewed and edited the content as needed and take full responsibility for the content of the published article.

## Data availability

The structure factors and atomic coordinates of full-length *Eh*APSK and *Eh*APSK_ΔKD_ determined by X-ray crystallography have been deposited in the Protein Data Bank under the accession codes 000023VK and 000023VL, respectively. In addition, the potential map of full-length *Eh*APSK obtained by single-particle cryo-EM has been deposited in the Electron Microscopy Data Bank under the accession code EMD-80056.

## Notes

### Competing Interest Statement

The authors have declared no competing interest.

## References

1. Schelle, M.W., Bertozzi, C.R., 2006. Sulfate metabolism in mycobacteria. Chembiochem. 7(10):1516–24.

2. Patron, N.J., Durnford, D.G., Kopriva, S., 2008. Sulfate assimilation in eukaryotes. Fusions, relocations, and lateral transfers. BMC Evol. Biol. 8:39.

3. Yi, H., Galant, A., Ravilious, G.E., et al., 2010. Sensing sulfur conditions. Simple to complex protein regulatory mechanisms in plant thiol metabolism. Mol. Plant 3(2):269–279.

4. Takahashi, H., Kopriva, S., Giordano, M., et al., 2011. Sulfur assimilation in photosynthetic organisms. Molecular functions and regulations of transporters and assimilatory enzymes. Annu. Rev. Plant Biol. 62:157–184.

5. Robbins, P.W., Lipmann, F., 1958. Separation of the two enzymatic phases in active sulfate synthesis. J. Biol. Chem. 233(3):681–5.

6. Coughtrie, M.W.H., 2016. Function and organization of the human cytosolic sulfotransferase (SULT) family. Chem. Biol. Interact. 259(Pt A):2–7.

7. Kusche, M., Oscarsson, L.G., Reynertson, R., et al., 1991. Biosynthesis of heparin. Enzymatic sulfation of pentasaccharides. J. Biol. Chem. 266(12):7400–9.

8. Suiko, M., Fernando, P.H., Sakakibara, Y., et al., 1992. Post-translational modification of protein by tyrosine sulfation: active sulfate PAPS is the essential substrate for this modification. Nucleic Acids Symp. Ser. (27):183–4.

9. Varin, L., DeLuca, V., Ibrahim, R.K., et al., 1992. Molecular characterization of two plant flavonol sulfotransferases. Proc. Natl. Acad. Sci. 89(4):1286–90.

10. Klaassen, C.D., Boles, J.W., 1997. Sulfation and sulfotransferases 5: the importance of 3’-phosphoadenosine 5’-phosphosulfate (PAPS) in the regulation of sulfation. FASEB J. 11(6):404–18.

11. van der Ploeg, J.R., Eichhorn, E., Leisinger, T., 2001. Sulfonate-sulfur metabolism and its regulation in *Escherichia coli*. Arch. Microbiol. 176(1-2):1–8.

12. Mendoza-Cózatl, D., Loza-Tavera, H., Hernandez-Navarro, A., et al., 2005. Sulfur assimilation and glutathione metabolism under cadmium stress in yeast, protists and plants. FEMS Microbiol. Rev. 29(4):653–71.

13. Hatzious, S.K., Bertozzi, C.R., 2011. The regulation of sulfur metabolism in *Mycobacterium tuberculosis*. PLOS Pathog. 7(7):e1002036.

14. Koprivova, A., Kopriva, S., 2014. Molecular mechanisms of regulation of sulfate assimilation: first steps on a long road. Front. Plant Sci. 5:589.

15. Rath, V.L., Verdugo, D., Hemmerich, S., 2004. Sulfotransferase structural biology and inhibitor discovery. Drug Discov. Today 9(23):1003–11.

16. Günal, S., Hardman, R., Kopriva, S., et al., 2019. Sulfation pathways from red to green. J. Biol. Chem. 294(33):12293–12312.

17. Mueller, J.W., Idkowiak, J., Gesteira, T.F., et al., 2018. Human DHEA sulfation requires direct interaction between PAPS synthase 2 and DHEA sulfotransferase SULT2A1. J. Biol. Chem. 293(25):9724–9735.

18. Lansdon, E.B., Fisher, A.J., Segel, I.H., 2004. Human 3’-phosphoadenosine 5’-phosphosulfate synthetase (isoform 1, brain): kinetic properties of the adenosine triphosphate sulfurylase and adenosine 5’-phosphosulfate kinase domains. Biochemistry 43(14):4356–65.

19. Brylski, O., Shrestha, P., House, P.J., et al., 2022. Disease-related protein variants of the highly conserved enzyme PAPSS2 show marginal stability and aggregation in cells. Front. Mol. Biosci. 9:860387.

20. Mueller, J.W., Shafqat, N., 2013. Adenosine-5′-phosphosulfate – a multifaceted modulator of bifunctional 3′-phospho-adenosine-5′-phosphosulfate synthases and related enzymes. FEBS J. 280(13):3050–7.

21. Mi-ichi, F., Yoshida, H., 2019. Unique Features of *Entamoeba* Sulfur Metabolism; Compartmentalization, Physiological Roles of Terminal Products, Evolution and Pharmaceutical Exploitation. Int. J. Mol. Sci. 20(19):4679.

22. Bradley, M.E., Rest, J.S., Li, W.H., et al., 2009. Sulfate Activation Enzymes: Phylogeny and Association with Pyrophosphatase. J. Mol. Evol. 68(1):1–13.

23. MacRae, I., Segel, I.H., 1997. ATP sulfurylase from filamentous fungi: which sulfonucleotide is the true allosteric effector? Arch. Biochem. Biophys. 337(1):17–26.

24. Ronosto, F., Martin, R.L., Segel, I.H., 1989. Sulfate-activating enzymes of *Penicillium chrysogenum*. The ATP sulfurylase.adenosine 5’-phosphosulfate complex does not serve as a substrate for adenosine 5’-phosphosulfate kinase. J. Biol. Chem. 264(16):9433–7.

25. MacRae, I.J., Segel, I.H., Fisher, A.J., 2001. Crystal Structure of ATP Sulfurylase from *Penicillium chrysogenum*:L Insights into the Allosteric Regulation of Sulfate Assimilation. Biochemistry 40(23):6795–804.

26. Lozano, R., Naghavi, M., Foreman, K., et al., 2012. Global and regional mortality from 235 causes of death for 20 age groups in 1990 and 2010: a systematic analysis for the Global Burden of Disease Study 2010. Lancet 380(9859):2095–128.

27. Quach, J., St-Pierre, J., Chadee, K., 2014. The future for vaccine development against *Entamoeba histolytica*. Hum. Vaccin. Immunother. 10(6):1514–1521.

28. Mi-ichi, F., Makiuchi, T., Furukawa, A., et al., 2011. Sulfate Activation in Mitosomes Plays an Important Role in the Proliferation of *Entamoeba histolytica*. PLOS Negl. Trop. Dis. 5(8):e1263.

29. Mi-ichi, F., Miyamoto, T., Takao, S., et al., 2015. *Entamoeba* mitosomes play an important role in encystation by association with cholesteryl sulfate synthesis. Proc. Natl. Acad. Sci. 112(22):E2884–90.

30. Mi-ichi, F., Miyamoto, T., Yoshida, H., 2017. Uniqueness of *Entamoeba* sulfur metabolism: sulfolipid metabolism that plays pleiotropic roles in the parasitic life cycle. Mol. Microbiol. 106(3):479–491.

31. Mi-ichi, F., Tsugawa, H., Arita, M., et al., 2022. Pleiotropic Roles of Cholesteryl Sulfate during Entamoeba Encystation: Involvement in Cell Rounding and Development of Membrane Impermeability. mSphere 7(4):e0029922.

32. Leyh, T.S., Taylor, J.C., Markham, G.D., 1988. The sulfate activation locus of *Escherichia coli* K12: cloning, genetic, and enzymatic characterization. J. Biol. Chem. 263(5):2409–16.

33. Besset, S., Vincourt, J.B., Amalric, F., et al., 2000. Nuclear localization of PAPS synthetase 1: a sulfate activation pathway in the nucleus of eukaryotic cells. FASEB J. 14(2):345–54.

34. Brunold, C., Suter, M., 1989. Localization of enzymes of assimilatory sulfate reduction in pea roots. Planta 179(2):228–34.

35. Bohrer, A.S., Yoshimoto, N., Sekiguchi, A., et al., 2015. Alternative translational initiation of ATP sulfurylase underlying dual localization of sulfate assimilation pathways in plastids and cytosol in *Arabidopsis thaliana*. Front. Plant Sci. 5:750.

36. Mi-ichi, F., Abu Yousuf, M., Nakada-Tsukui, K., et al., 2009. Mitosomes in *Entamoeba histolytica* contain a sulfate activation pathway. Proc. Natl. Acad. Sci. 106(51):21731–6.

37. Santos, H.J., Nozaki, T., 2022. The mitosome of the anaerobic parasitic protist *Entamoeba histolytica*: A peculiar and minimalist mitochondrion-related organelle. J. Eukaryot. Microbiol. 69(6):e12923.

38. Mi-ichi, F., Nozawa, A., Yoshida, H., et al., 2015. Evidence that the *Entamoeba histolytica* Mitochondrial Carrier Family Links Mitosomal and Cytosolic Pathways through Exchange of 3’-Phosphoadenosine 5’-Phosphosulfate and ATP. Eukaryot. Cell 14(11):1144–50.

39. Mi-ichi, F., Ishikawa, T., Tam, V.K., et al., 2019. Characterization of *Entamoeba histolytica* adenosine 5’-phosphosulfate (APS) kinase; validation as a target and provision of leads for the development of new drugs against amoebiasis. PLOS Negl. Trop. Dis. 13(8):e0007633.

40. Gay, S.C., Fribourgh, J.L., Donohoue, P.D., et al., 2009. Kinetic properties of ATP sulfurylase and APS kinase from *Thiobacillus denitrificans*. Arch. Biochem. Biophys. 489(1-2):110–7.

41. Gay, S.C., Segel, I.H., Fisher, A.J., 2009. Structure of the two-domain hexameric APS kinase from *Thiobacillus denitrificans*: structural basis for the absence of ATP sulfurylase activity. Acta Crystallogr. D Biol. Crystallogr. 65(Pt 10):1021–31.

42. Yu, Z., Lansdon, E.B., Segel, I.H., et al., 2007. Crystal structure of the bifunctional ATP sulfurylase-APS kinase from the chemolithotrophic thermophile *Aquifex aeolicus*. J. Mol. Biol. 365(3):732–43.

43. Ravilious, G.E., Nguyen, A., Francois, J.A., et al., 2012. Structural basis and evolution of redox regulation in plant adenosine-5’-phosphosulfate kinase. Proc. Natl. Acad. Sci. 109(1):309–14.

44. Lansdon, E.B., Segel, I.H., Fisher, A.J., 2002. Ligand-induced structural changes in adenosine 5’-phosphosulfate kinase from *Penicillium chrysogenum*. Biochemistry 41(46):13672–80.

45. Herrmann, J., Nathin, D., Lee, S.G., 2015. Recapitulating the Structural Evolution of Redox Regulation in Adenosine 5’-Phosphosulfate Kinase from Cyanobacteria to Plants. J. Biol. Chem. 290(41):24705–14.

46. Harjes, S., Bayer, P., Scheidig, A.J., 2005. The Crystal Structure of Human PAPS Synthetase 1 Reveals Asymmetry in Substrate Binding. J. Mol. Biol. 290(41):24705–14.

47. Sekulic, N., Dietrich, K., Paarmann, I., et al., 2007. Elucidation of the active conformation of the APS-kinase domain of human PAPS synthetase 1. J. Mol. Biol. 367(2):488–500.

48. MacRae, I.J., Segel, I.H., Fisher, A.J., 2000. Crystal structure of adenosine 5’-phosphosulfate kinase from *Penicillium chrysogenum*. Biochemistry 39(7):1613–21.

49. Punjani, A., Rubinstein, J.L., Fleet, D.J., et al., 2017. cryoSPARC: algorithms for rapid unsupervised cryo-EM structure determination. Nat. Methods 14(3):290–296.

50. Jumper, J., Evans, R., Pritzel, A., et al., 2021. Highly accurate protein structure prediction with AlphaFold. Nature 596(7873):583–589.

51. Dolezal, P., Dagley, M.J., Kono, M., et al., 2010. The essentials of protein import in the degenerate mitochondrion of *Entamoeba histolytica*. PlOS Pathog. 6(3):e1000812.

52. Liu, A.Y., Koga, H., Goya, C., et al., 2023. Quick and affordable DNA cloning by reconstitution of Seamless Ligation Cloning Extract using defined factors. Genes Cells 28(8):553–562.

53. Kabsch, W., 2010. XDS. Acta Crystallogr. D Biol. Crystallogr. 66(Pt 2):125–132.

54. Skubák, P., Araç, D., Bowler, M.R., et al., 2018. A new MR-SAD algorithm for the automatic building of protein models from low-resolution X-ray data and a poor starting model. IUCrJ 5(Pt 2):166–171.

55. Collaborative Computational Project, Number 4, 1994. The CCP4 suite: programs for protein crystallography. Acta Crystallogr. D Biol. Crystallogr. 50(Pt 5):760–763.

56. Cowtan, K., 2006. The Buccaneer software for automated model building. 1. Tracing protein chains. Acta Crystallogr. D Biol. Crystallogr. 62(Pt 9):1002–11.

57. Cowtan, K., 2008. Fitting molecular fragments into electron density. Acta Crystallogr. D Biol. Crystallogr. 64(Pt 1):83–9.

58. Vagin, A., Teplyakov, A., 1997. MOLREP: an Automated Program for Molecular Replacement. J. Appl. Cryst. 30:1022–1025.

59. McCoy, A.J., Grosse-Kunstleve, R.W., Adams, P.D., et al., 2007. Phaser crystallographic software. J. Appl. Cryst. 40(Pt 4):658–674.

60. Murshudov, G.N., Vagin, A.A., Dodson, E.J., 1997. Refinement of macromolecular structures by the maximum-likelihood method. Acta Crystallogr. D Biol. Crystallogr. 53(Pt 3):240–255.

61. Emsley, P., Cowtan, K., 2004. Coot: model-building tools for molecular graphics. Acta Crystallogr. D Biol. Crystallogr. 60(Pt 12 Pt 1):2126–2132.

62. DeLano, W.L., 2002. The PyMOL Molecular Graphics System. http://www.pymol.org

63. Pettersen, E.F., Goddard, T.D., Huang, C.C., et al., 2021. UCSF ChimeraX: Structure visualization for researchers, educators, and developers. Protein Sci. 30(1):70–82.

64. Mastronarde, D.N., 2005. Automated electron microscope tomography using robust prediction of specimen movements. J. Struct. Biol. 152(1):36–51.

65. Bepler, T., Morin, A., Rapp, M., et al., 2019. Positive-unlabeled convolutional neural networks for particle picking in cryo-electron micrographs. Nat. Methods 16(11):1153–1160.

