## Supplementary Figures and Tables for "Structural insights into interdomain interactions in *Entamoeba histolytica* APS kinase"

7 figures and 4 tables.

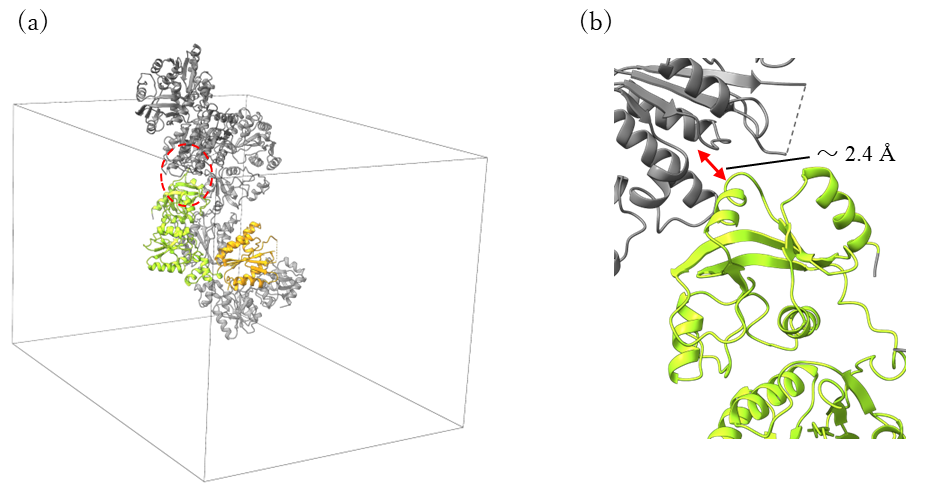

**Figure S1. Asymmetric unit of wild-type *Eh*APSK.** (a) Asymmetric unit of *Eh*APSK revealed by X-ray crystallography. A cube enclosing the tetramer represents the unit cell (gray cube). A tetramer is formed by two W-shaped dimers arranged in a half-shifted and face-to-face orientation. (b) The tip of SLD is in close proximity (~2.4 Å) to the APS-binding site of KD in the other dimer.

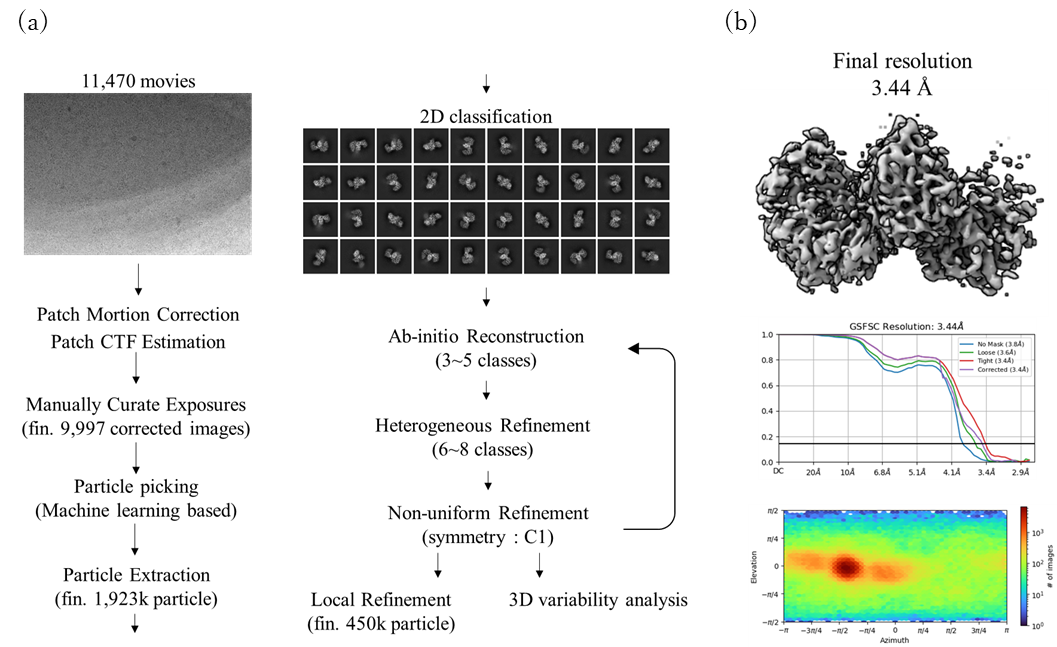

**Figure S2. Workflow of the single-particle cryo-EM.** (a) Single-particle cryo-EM was performed using cryoSPARC as shown in the workflow. (b) The final map, the Gold-Standard FSC curve, and the angular distribution plot obtained by local refinement are shown. Multiple masking regions were tested for local refinement and 3D classification; however, no improvement of the map beyond that obtained by local refinement was achieved. Also, due to the low resolution of the map, atomic model building was not performed.

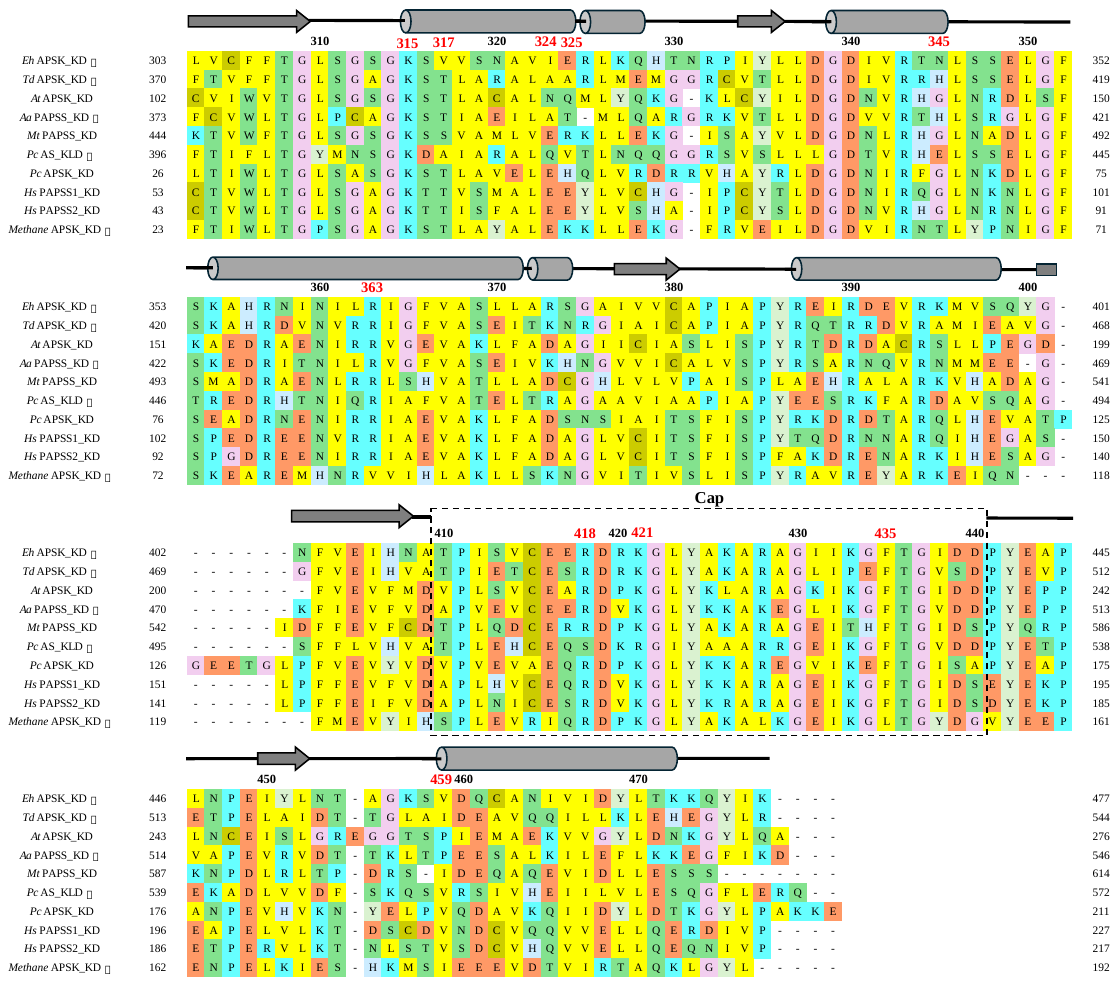

**Figure S3.** **Multiple sequence alignment of the KD region of *Eh*APSK.** Residues are color-coded according to their physicochemical properties. The numbering above the sequence corresponds to the residue numbering of *Eh*APSK, and residues discussed in the main text are indicated by red. Cylinders and arrows represent the α-helices and β-sheets of *Eh*APSK, respectively, and the unmodeled cap region is indicated by a dashed box. Asterisks (*) indicate the “closed” KD dimer for each sequence.

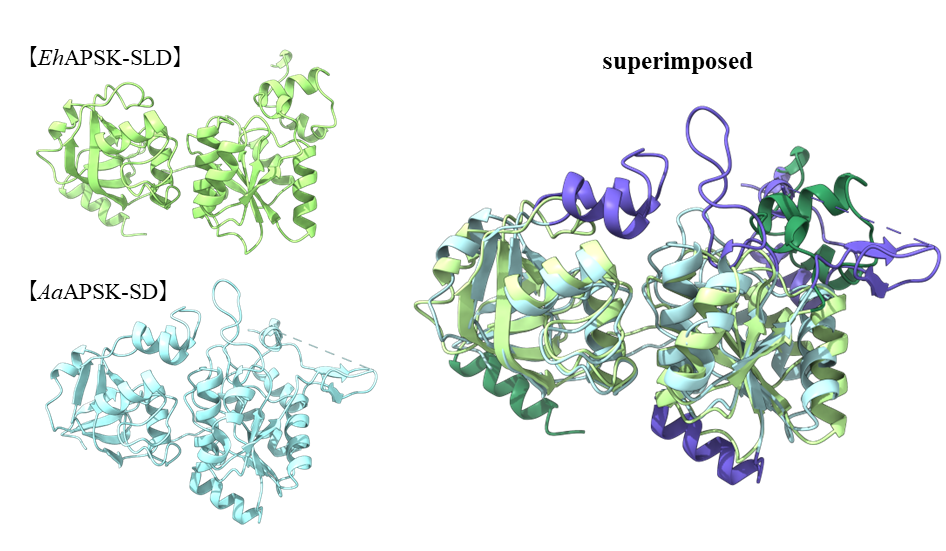

**Figure S4. Unique structure of SLD.** Structural superposition of the SLD of *Eh*APSK (green) and the SD of *Aa*PAPSS(PDB ID: 2GKS) (blue). Regions showing structurally different are highlighted in darker colors.

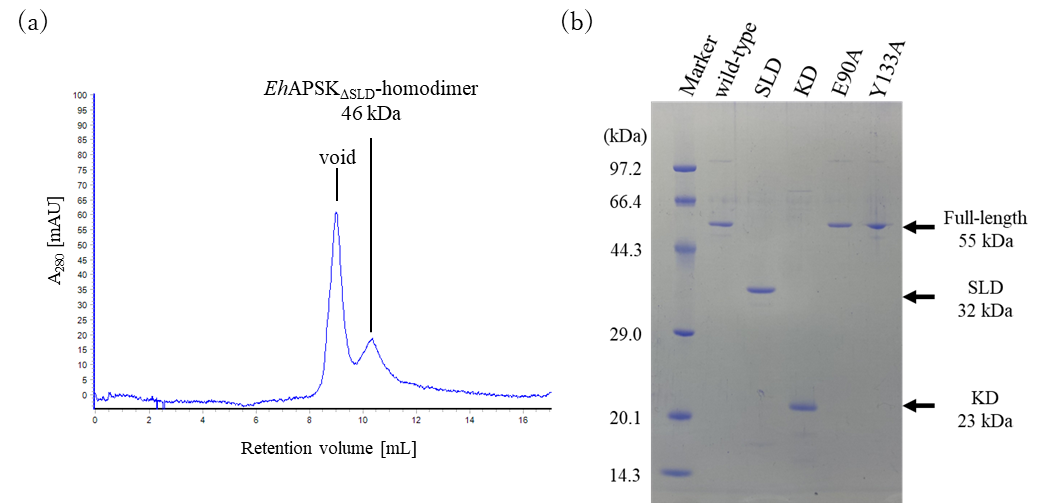

**Figure S5. Analysis of purified samples** (a) SEC profile of *Eh*APSK_ΔSLD_ after SUMO tag removal. *Eh*APSK_ΔSLD_ was structurally unstable and highly prone to aggregation after removal of the SUMO tag, with most aggregated *Eh*APSK_ΔSLD_ eluting in the void volume. In subsequent experiments, the low-concentration dimer fraction separated by SEC was used directly. (b) SDS-PAGE analysis of each purified sample. Samples with sufficient purity were used for subsequent structural analysis and activity measurements.

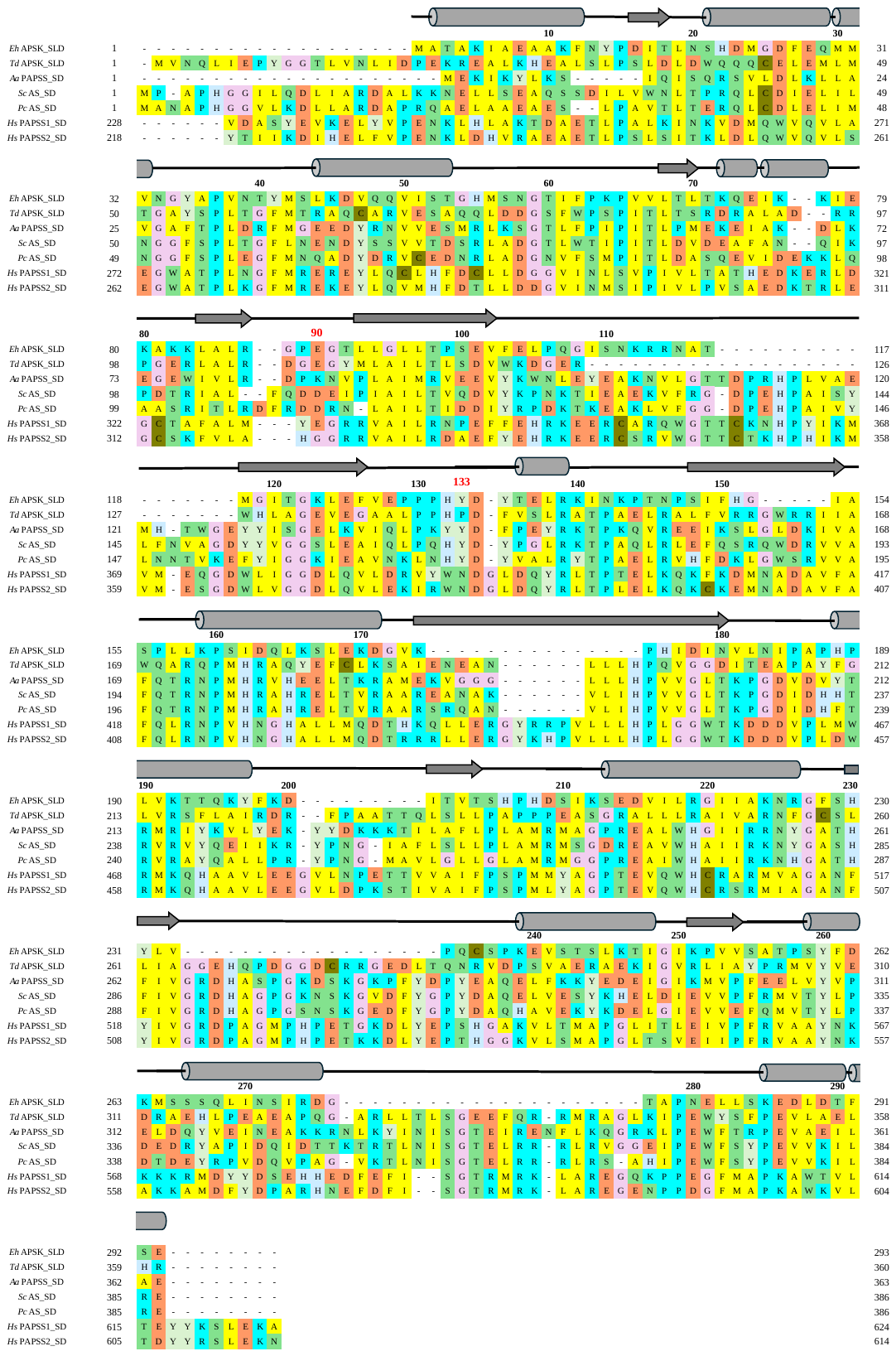

**Figure S6.** **Multiple sequence alignment of the SLD region of *Eh*APSK.** Residues are color-coded according to their physicochemical properties. The numbering above the sequence corresponds to the residue numbering of *Eh*APSK, and residues discussed in the main text are indicated by red. Cylinders and arrows represent the α-helices and β-sheets of *Eh*APSK.

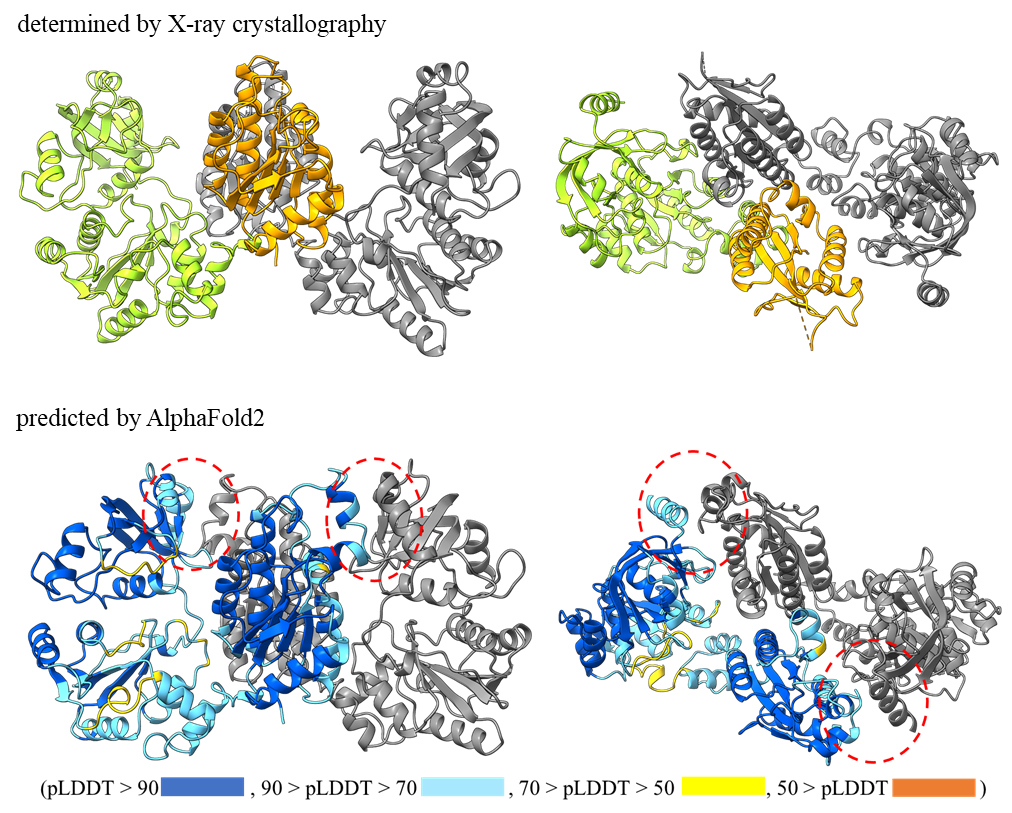

**Figure S7. Comparison between the crystal structure and the AlphaFold2-predicted structure of *Eh*APSK.** The crystal structure of wild-type *Eh*APSK is shown in the upper row (A chain is colored with green and orange), and the AlphaFold2-predicted structure of wild-type *Eh*APSK is shown in the lower row (A chain is colored by pLDDT: Dark blue 90-100, light blue 70-90, yellow 50-70, and orange 0-50). The structures of each domain are similar (RMSD_SLD = 1.031 [224 pruned atoms] (2.297 [all 297 pairs]); RMSD_KD = 0.785 [139 pruned atoms] (2.095 [all 149 pairs])), but the overall protomer alignment is poor (RMSD_full-lemgth = 0.673 [134 pruned atoms] (5.275 [all 452 pairs])). This difference in relative domain positions may explain why molecular replacement using the predicted structure was unsuccessful in the crystallographic analysis. In the predicted structure, SLD approaches KD, and the tip of SLD interacts with the cap region of KD (red dotted circles).

**Table S1. Data statistics for X-ray crystallographic analysis.**

|  | Full-length *Eh*APSK | | *Eh*APSK_ΔKD_ |
| --- | --- | --- | --- |
|  | SAD |  |  |
| PDB ID |  | 000023VK | 000023VL |
| Space group | *C*222_1_ | *C*222_1_ | *P*2_1_2_1_2_1_ |
| *a, b, c*（Å^2^） | 146.7, 158.3, 227.7 | 144.4, 158.4, 226.2 | 53.6, 55.7, 105.8 |
| *α, β, γ*（°） | 90.0, 90.0, 90.0 | 90.0, 90.0, 90.0 | 90.0, 90.0, 90.0 |
| Beam line | SPring-8, BL44XU | SPring-8, BL44XU | SPring-8, BL44XU |
| Wavelength（Å^2^） | 0.9000 | 0.9000 | 0.9000 |
| Resolution（Å^2^） | 50-2.90 (3.07-2.90) | 50-2.59 (2.75-2.59) | 50-2.10 (2.22-2.10) |
| *R*_merge_ | 0.312 (6.596) | 0.066 (2.391) | 0.127 (1.503) |
| *I*/σ (*I*) | 9.6 (0.6) | 14.36 (0.80) | 12.95 (1.99) |
| Completeness (%) | 99.5 (97.4) | 99.5 (98.1) | 98.5 (96.4) |
| Redundancy | 14.1 (14.1) | 6.1 (6.2) | 6.1 (6.4) |
| Refinement |  |  |  |
| Resolution (Å^2^) |  | 19.97-2.60 | 19.93-2.10 |
| *R*_work_/*R*_free_ |  | 0.227/0.265 | 0.226/0.286 |
| average *B*-factor (Å^2^) |  | 111.3 | 54.8 |

Values in parentheses indicate the highest-resolution shell. Additionally, 5% of the total reflections were used for *R*_free_ calculation.

**Table S2. Domain comparison between *Eh*APSK and AS/APSK from other organisms.**

(a)

| SD / SLD of | Protomer Structure  (N terminal / C terminal) | Sequance Identity with SLD_M1~E293 of *Eh*APSK  (%) | RMSD (Å) |
| --- | --- | --- | --- |
| *Aa*PAPSS_M1~E363 （uniprot：O67174、PDB ID：2GKS） | **SD** / KD (or KLD) | 22.58 | 1.148 [133 pruned atom]  (6.632 [all 281 pairs]) |
| *Pc*AS_M1~E386 （uniprot：Q12650、PDB ID：1I2D） | **SD** / KLD | 22.46 | 1.103 [121 pruned atom]  (5.803 [all 291 pairs]) |
| *Td*APSK_M1~R360 （uniprot：Q3SM86、PDB ID：3CR8） | **SLD** / KD | 20.15 | 1.190 [104 pruned atom]   (5.353 [all 280 pairs]) |
| *Sc*AS_M1~E386 （uniprot：P08536、PDB ID：1G8F） | **SD** / KLD | 19.93 | 0.953 [121 pruned atom]  (5.854 [all 293 pairs]) |
| *Hs*PAPSS1_V228~A624 （uniprot：O43252、PDB ID：1X6V） | KD / **SD** | 18.62 | 1.322 [106 pruned atom]   (6.067 [all 291 pairs]) |
| *Hs*PAPSS2_Y218~N614 （uniprot：O95340） | KD / **SD** | 14.83 | ― |

(b)

| KD / KLD of | Protomer Structure  (N terminal / C terminal) | Sequance Identity with KD_L303~K477 of *Eh*APSK  (%) | RMSD (Å) |
| --- | --- | --- | --- |
| *Td*APSK_F370~R544 （uniprot：Q3SM86、PDB ID：3CR8） | SLD / **KD** | 54.86 | 0.856 [133 pruned atom] (1.391 [all 146 pairs]) |
| *At*APSK_M1~A276 （uniprot：Q43295、PDB ID：3UIE） | KD | 49.71 | 0.986 [122 pruned atom]  (2.730 [all 147 pairs]) |
| *Aa*PAPSS_F373~D546 （uniprot：O67174、PDB ID：2GKS） | SD / **KD (or KLD)** | 46.24 | 0.857 [118 pruned atom]  (3.185 [all 147 pairs]) |
| *Mt*APSK_K444~S614 （uniprot：A5U1Y4、PDB ID：4BZP） | GTPase / **KD** | 45.29 | 0.945 [116 pruned atom]  (2.447 [all 142 pairs]) |
| *Pc*AS_F396~Q572 （uniprot：Q12650、PDB ID：1I2D） | SD / **KLD** | 45.14 | 0.934 [118 pruned atom]  (2.510 [all 149 pairs]) |
| *Pc*APSK_M1~E211 （uniprot：Q12657、PDB ID：1M7G） | KD | 44.57 | 0.968 [121 pruned atom]  (3.186 [all 147 pairs]) |
| *Hs*PAPSS1_M1~P227 （uniprot：O43252、PDB ID：1X6V） | **KD** / SD | 42.53 | 0.939 [130 pruned atom]  (1.960 [all 144 pairs]) |
| *Hs*PAPSS2_M1~P217 （uniprot：O95340、PDB ID：8I1O） | **KD** / SD | 42.53 | 1.059 [125 pruned atom]  (3.224 [all 148 pairs]) |
| *Methane*APSK_G1~L192 （uniprot：A0AA82WPC2、PDB ID：8A8H） | KD | 37.65 | 1.042 [109 pruned atom]  (1.834 [all 140 pairs]) |

The amino acid sequence identity and RMSD values for (a) SD/SLD and (b) KD/KLD are summarized in the table above.

**Table S3. Primer sequences used for mutant construction.**

| name | Tm (℃) | sequence |
| --- | --- | --- |
| *Eh*APSK_ΔKD__forward | 60.68 | 5’ - TAGTGAAATTTGACTGCAGTCTAGATAGGTAATCTC -3’ |
| *Eh*APSK_ΔKD__reverse | 59.75 | 5’ - ACTGCAGTCAAATTTCACTAAAAGTATCCAAATCTTC -3’ |
| *Eh*APSK_ΔSLD__KD_forward | 61.49 | 5’ - TGGTGGTAGCTATCCACCAACTGATAAACAAGG - 3’ |
| *Eh*APSK_ΔSLD__KD_reverse | 61.43 | 5’ - CTTGTCTTCATTTAATGTATTGCTTCTTTGTGAGATAG - 3’ |
| *Eh*APSK_ΔSLD__pET-SUMO_forward | 62.29 | 5’ - ATACATTAAATGAAGACAAGCTTAGGTATTTATTCG - 3’ |
| *Eh*APSK_ΔSLD__pET SUMO_reverse | 62.56 | 5’ - TTGGTGGATAGCTACCACCAATCTGTTCTCTG - 3’ |
| E90A_forward | 59.25 | 5’ - GAGGTCCAGCAGGTACTTTAC - 3 |
| E90A_reverse | 59.88 | 5’ - AAGTACCTGCTGGACCTCTC - 3’ |
| Y133A_forward | 61.85 | 5’ - CACCTCATGCCGATTATACTG - 3’ |
| Y133A_reverse | 61.69 | 5’ - GTATAATCGGCATGAGGTGG - 3’ |

Primers were designed to have melting temperature (Tm) of approximately 60℃.

**Table S4. Statistic table for cryo-EM analysis of *Eh*APSK.**

| EMDB ID | EMD-80056 |
| --- | --- |
| Magnification | 105,000 |
| Voltage (kV) | 300 |
| Total dose (e^-^/Å^2^) | 65 |
| Defocus range (µm) | -0.8 to -1.8 |
| Pixel size (Å) | 0.675 |
| Collected spherical aberration | 0.066 |
| Symmetry imposed | C1 |
| # of movies | 11,470 |
| # of Initial particles | 1,923,378 |
| # of Final particle | 450,150 |
| Resolution (Å) GSFSC = 0.143 | 3.44 |

The imaging conditions using the Titan Krios and the final statistics of the analyzed dataset are summarized in the table above.
